# Ventral tegmental area glutamate neurons mediate the nonassociative consequences of traumatic stress

**DOI:** 10.1101/2021.04.02.438264

**Authors:** Dillon J. McGovern, Koy L. Ecton, David T. Huynh, Andrew R. Rau, Shane T. Hentges, Annie Ly, Michael V. Baratta, David H. Root

## Abstract

Exposure to trauma is a risk factor for the development of a number of mood disorders, and may enhance vulnerability to future adverse life events. Recent data implicate ventral tegmental area (VTA) glutamate neuronal activity as functionally important for signaling aversive or threating stimuli. However, it is unknown whether VTA glutamate neurons regulate transsituational outcomes that result from stress and whether these neurons are sensitive to stressor controllability. This work established an operant mouse paradigm to examine the impact of stressor controllability on VTA glutamate neuron function and stressor outcome. Uncontrollable (inescapable) stress, but not physically identical controllable (escapable) stress, produced social avoidance in male mice. Cell-type-specific calcium recordings showed that both controllable and uncontrollable stressors increased VTA glutamate neuronal activity. Chemogenetic reduction of VTA glutamate neuron activity prevented the behavioral sequelae of uncontrollable stress. Our results provide causal evidence that mice can be used to model stressor controllability and that VTA glutamate neurons may contribute to transsituational stressor outcomes, such as social avoidance and exaggerated fear that are observed within trauma-related disorders.

## INTRODUCTION

Exposure to traumatic life events has been implicated in a number of mood and anxiety disorders. Traumatic experiences can lead to a broad range of negative consequences that are supported by both associative (e.g., avoidance of or distress due to stimuli related to the trauma) and nonassociative (e.g., exaggerated transsituational responding to circumstances unrelated to the trauma) processes (Der-Avakian et al., 2014; Lissek and Grillon, 2012; Lissek and van Meurs, 2015). Although exposure-based therapies are considered an efficacious intervention for a variety of stress-linked disorders, a substantial number of individuals fail to achieve long-term symptom reduction (Kline et al., 2018; Morina et al., 2014; Watts et al., 2013). Treatment resistance, in part, may be due to the fact that exposure therapy primarily targets the learned, associative components of the disorder, leaving the nonassociative aspects inadequately addressed (Markowitz and Fanselow, 2020).

Well known for its role in reward, the ventral tegmental area (VTA) also plays an important role in aversion (Lammel et al., 2014). The VTA is cellularly heterogeneous mid brain region with diverse molecular properties. In addition to dopamine neurons, the VTA contains GABAergic and glutamatergic subpopulations (Mingote et al., 2017; Morales and Root, 2014) that exhibit distinct synaptic architecture and connectivity (Lammel et al., 2014; Morales and Margolis, 2017). Subpopulations of dopamine, GABA, and glutamate neurons have been shown to participate in aversive processing (Lammel et al., 2012; Root et al., 2014; Tan et al., 2012). Notably, VTA glutamate neurons are also activated by defensive responses to perceived threat, such as initiating escape from looming or predator odor threats, or maintaining mobility during the forced swim test (Barbano et al., 2020). Even in the absence of an external threat, photostimulation of VTA VGluT2 projections to either the nucleus accumbens shell or lateral habenula is sufficient to form associative phenomena such as conditioned place aversion (Qi et al., 2016; Root et al., 2014). Based on results showing the involvement of VTA glutamate neurons in aversion or threat, we hypothesized that VTA glutamatergic neurons contribute to the transsituational behavioral consequences that follow stress.

In the present experiments, we adapted a stressor controllability paradigm from rats to mice in order to examine the role of VTA glutamate neurons in mediating the transsituational consequences of inescapable stress. Prior work, largely in rats, has shown that exposure to uncontrollable stress produces behavioral sequelae (e.g. reduced social exploration, exaggerated fear) that do not occur with physically-identical controllable stress (Maier and Watkins, 2005). Further, uncontrollable stress alters later behavior in circumstances very different from the original stress experience, demonstrating that outcomes dependent on the uncontrollability of the stressor are mediated by a nonassociative process (Greenwood et al., 2010; Landgraf et al., 2015; Maier and Watkins, 2005). We found in male mice that inescapable tailshock stress (IS) produced social exploration deficits that were absent if the tailshock was escapable (ES). Calcium imaging showed that tailshock robustly increased the activity of VTA glutamate neurons independent of shock controllability. Chemogenetic inhibition of VTA glutamate neurons during tailshock blocked IS-induced social exploration deficits and exaggerated footshock-elicited freezing responses. Overall, we interpret these data to reflect that stress-induced activation of VTA glutamate neurons is required for the development of the nonassociative behavioral consequences that follow uncontrollable stress. Further, these experiments establish a mouse model of stressor controllability that can be combined with other genetically-diverse mouse driver lines for identifying the cell types and circuits underlying the nonassociative aspects of traumatic stress exposure.

## RESULTS

### Stress-induced Social Avoidance Depends on the Uncontrollability of the Stressor in Male Mice

We first determined if deficits in social behavior, a common feature of anxiety disorders (Kessler et al., 1999; Shin and Liberzon, 2010), would result from prior exposure to inescapable stress (IS). Male and female C57BL/6J mice received IS or no stress (homecage controls; HC) and twenty-four hours later sociability was evaluated with the three-chamber preference test (Figure 1A). The 10-min social preference test was conducted in a novel apparatus and in a novel procedure room so as to minimize the presence of common cues between the stress and behavioral testing environments. A mixed ANOVA yielded a significant Sex X Stimulus interaction, F(1,14) = 5.5, p < 0.05. Sidak-adjusted pairwise comparisons indicated that IS males spent significantly less time with the social stimulus than no stress HC males, p < 0.05 (**Figure 1B**). In addition, no stress HC males demonstrated a significant preference for social stimulus interaction (p < 0.001), while IS males showed no difference in time spent between the social and non-social stimuli (Figure 1B). In contrast to males, both HC and IS females had a preference for spending time with a novel conspecific versus exploring an empty chamber (p < 0.05) (Figure 1C). These results establish that prior inescapable tailshock induces social avoidance in male mice.

**Figure 1:**
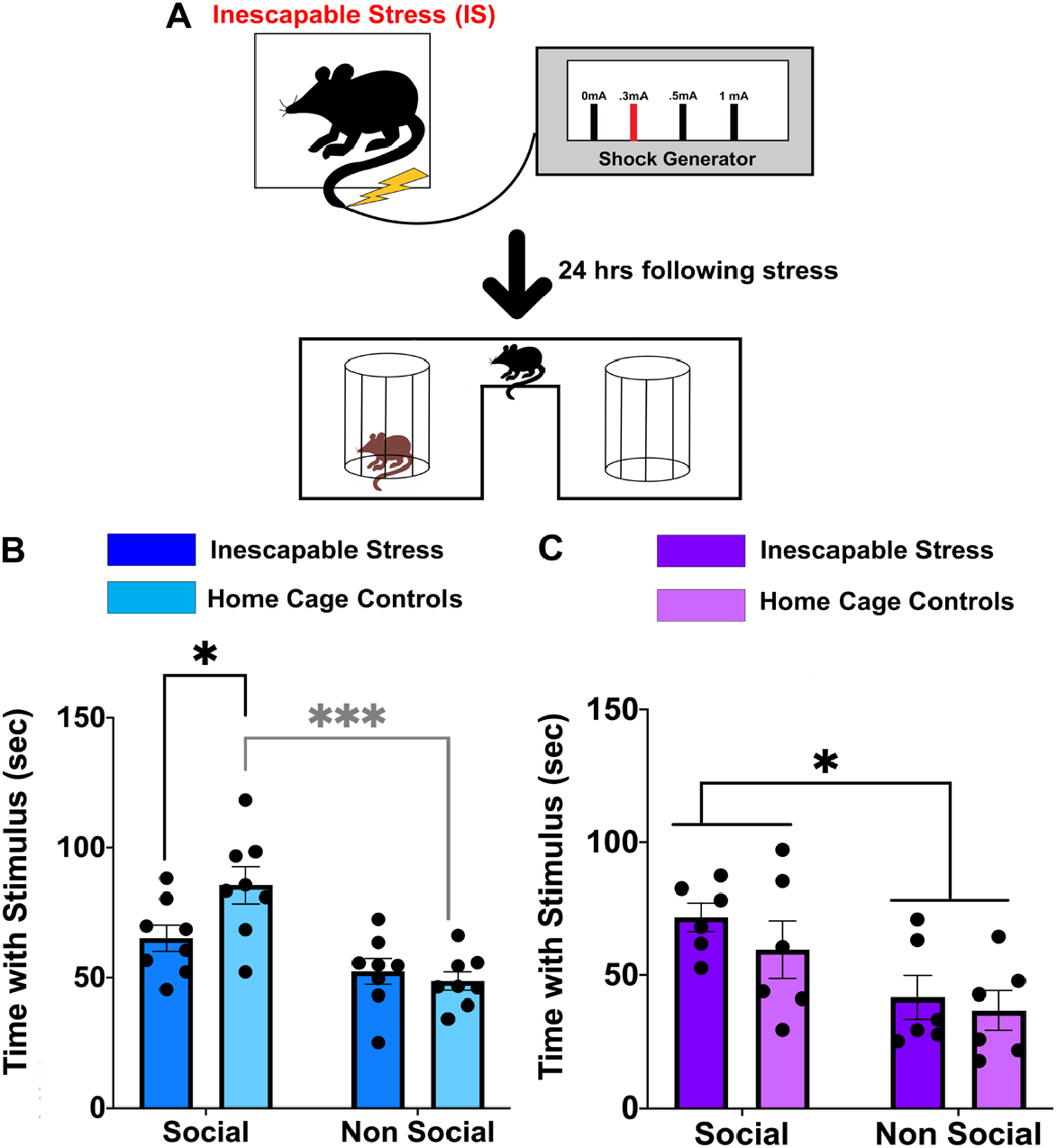
Inescapable stress reduces social exploration in male, but not female, C57BL/6J mice. (A) Schematic of the experimental timeline. Mice first received either inescapable stress (IS; 100 tailshocks, 5 sec duration each) or home cage treatment (no tailshock) followed by testing in the three-chamber sociability paradigm 24 hr later. (B-C) Quantification of interaction behavior with a social (same-sex conspecific) or a non social (empty chamber) stimulus for (B) male and (C) female subjects. Data are mean ± SEM. * p < 0.05, *** p < 0.001; ANOVA with Sidak’s multiple comparison post hoc test.

We next tested whether the social-related consequence of tailshock was dependent on its uncontrollability. Here, pairs of male mice received physically identical tailshocks, with one member having control over shock termination (Escapable stress; ES), and the other yoked to the ES subject but having no control (yoked inescapable stress; Y-IS). For each trial, tailshock was terminated for both subjects when the ES mouse completed the wheel-turn instrumental response criterion. Thus, both ES and Y-IS mice received exactly the same duration, intensity, and temporal pattern of tailshocks during stress treatment (**Figure 2A**). ES mice acquired the wheel-turn controlling response, significantly increasing the number of required quarter wheel-turns across the session until reaching the maximum response criterion (12 quarter-wheel turns), Friedman’s test X2(9) = 30.941, p < 0.001 (Figure 2A). Sidak-adjusted pairwise comparisons showed that the criterion quarter wheel-turns completed during the initial block of ten tailshock trials was significantly less than the last five blocks of ten tailshock trials, all p < 0.05. There was no significant change in latency to complete the criterion of quarter wheel-turns to terminate tailshocks over the training session, Χ2(9) = 11.69, p > 0.05.

**Figure 2:**
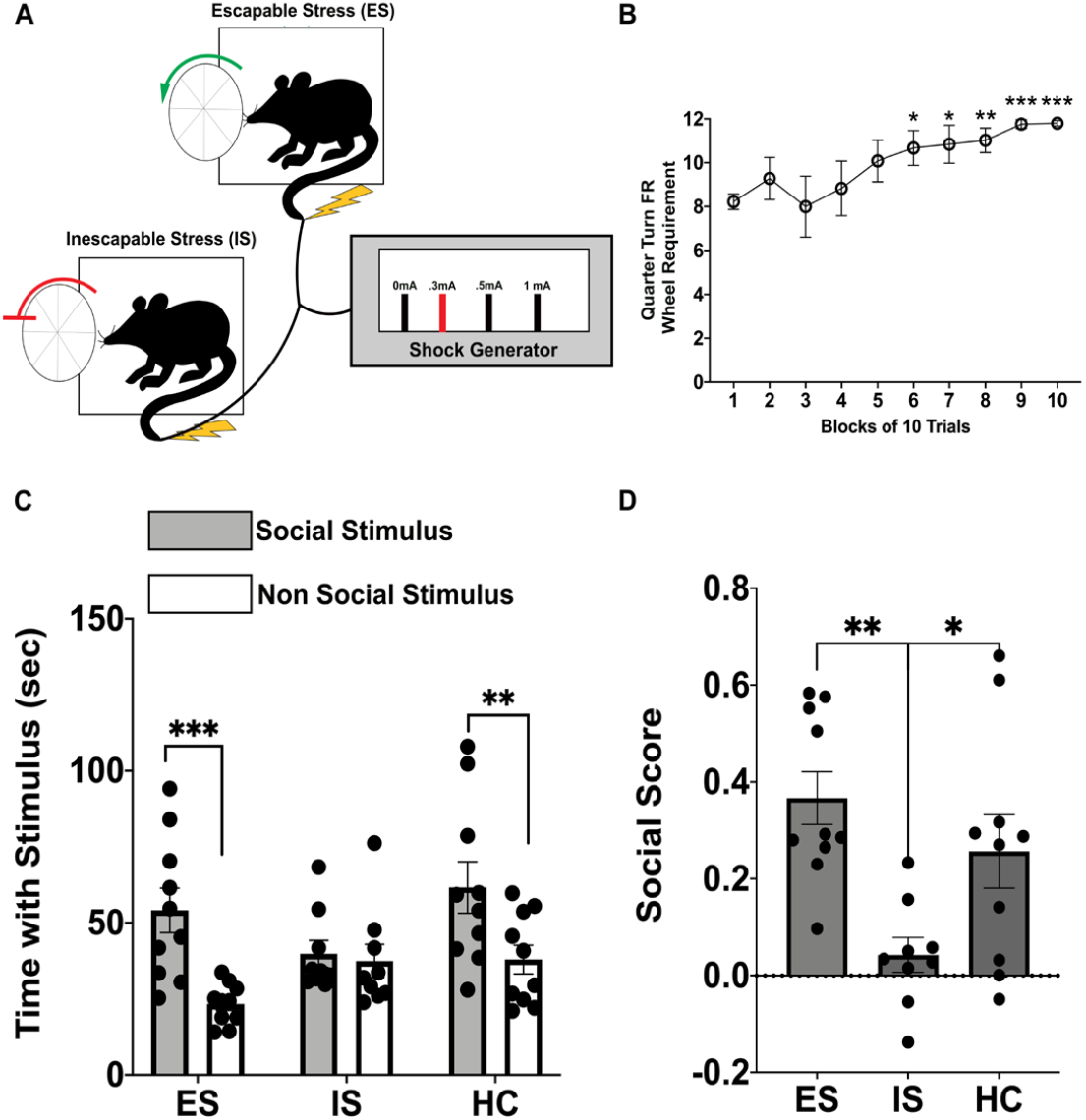
Stress-induced reduction in social exploration is depedent on the controllability of the stressor in male mice. (A) Illustration of the stressor controllability procedure. Escapable (ES), unlocked wheel and yoked inescapable stress (IS), locked wheel, male mice received 100 identical tailshocks (average intertrial interval 60 sec) that were terminated by the ES mouse. (B) Number of quarter turns of the wheel attained as the escape requirement in blocks of 10 trials. (C) Quantification of interaction behavior with a social (same-sex conspecific) or a non social (empty chamber) stimulus. ES and home cage (HC) subjects spent more time with the social stimulus than the non social stimulus. IS mice spent equal time with both stimulus types. (D) Preference for the social stimulus compared to the non social stimulus (social score) was significantly different between ES and IS groups. Data are mean ± SEM. * p < 0.05, ** p < 0.01, *** p < 0.001; ANOVA with Sidak’s multiple comparison post hoc test.

One day following stress treatment, ES, Y-IS, and no stress HC mice were tested in the three-chamber socialbility paradigm. The mixed ANOVA yielded a Group x Stimulus interaction, F(2,26) = 3.56, p < 0.05. Sidak-adjusted pairwise comparisons indicated that both ES and HC mice spent significantly more time with the social stimulus than the non-social stimulus (ES: p < 0.001; HC: p < 0.01), however preference for the conspecific was absent in Y-IS mice (p > 0.05) (Figure 2B). Social preference was quantified using a social score that normalized the time spent with the social stimulus relative to the time spent with the non-social stimulus.

ES mice had a significantly higher social score than Y-IS mice, main effect of Group: F(2,26) = 6.27, p < 0.01, Sidak-adjusted pairwise comparison p < 0.01. Importantly, the social score of Y-IS mice was near 0, indicating a complete lack of preference between social and non-social stimuli (**Figure 2C**). Total social interaction time was significantly less in IS mice when compared to home cage controls using an unpaired t-test, t(17)=2.457, p=0.0342, reconfirming that inescapable stress is sufficient to impair social exploration. These results establish that stressor-induced reduction in social exploration with a novel conspecific is controllability dependent, with Y-IS reducing social exploration but not physically identical ES.

### VTA Glutamate Neurons Respond to Both Uncontrollable and Controllable Stressors

In addition to dopamine and GABA neurons, the VTA contains glutamate-releasing VGluT2-expressing neurons (Morales and Root, 2014). VTA VGluT2 neurons show increased neuronal activity following a wide-range of aversive events (Root et al., 2018), including shock (Root et al., 2020), and can produce place aversion following their photostimulation in a projection-defined manner (Qi et al., 2016; Root et al., 2014; Wang et al., 2015). To understand whether controllability of the stressor modifies the activation of VTA glutamate neurons, we implanted an optical fiber into the VTA to simultaneously record changes in GCaMP6m fluorescence of VTA VGluT2 neurons in male VGluT2-IRES::Cre mice during the controllability paradigm (Figure 3A). VGluT2-IRES::Cre mice were randomly assigned to receive either escapable shock (ES) or physically identical inescapable tailshock (Y-IS). First, we found that Ca2+ activity of VTA VGluT2 neurons significantly increased immediately following tailshock when compared to pre-stimulus baselines for both experimental conditions during both the early phase acquisition of the wheel-turn response (initial 20 trials) and late phase (final 20 trials) of the session. Wilcoxon sign ranked tests indicate that Ca2+ activity was significantly higher at shock than baseline during ES early phase Z=60.712, p <0.05, IS early phase Z=49.848, p <0.05, ES late phase Z=23.728, p<0.05, and IS late phase Z=8.524, p<0.05. Ca2+ activity was not significantly different between ES and IS conditions at shock onset for the early stage t(10)=0.4439, p=0.666 or for the late stage t(10)=0.6476, p=0.5318 (Figure 3B-C).Thus, the strong activation of VTA glutamate neurons to tailshock was not sensitive to its controllability across the 100-trial tailshock session.

**Figure 3.**
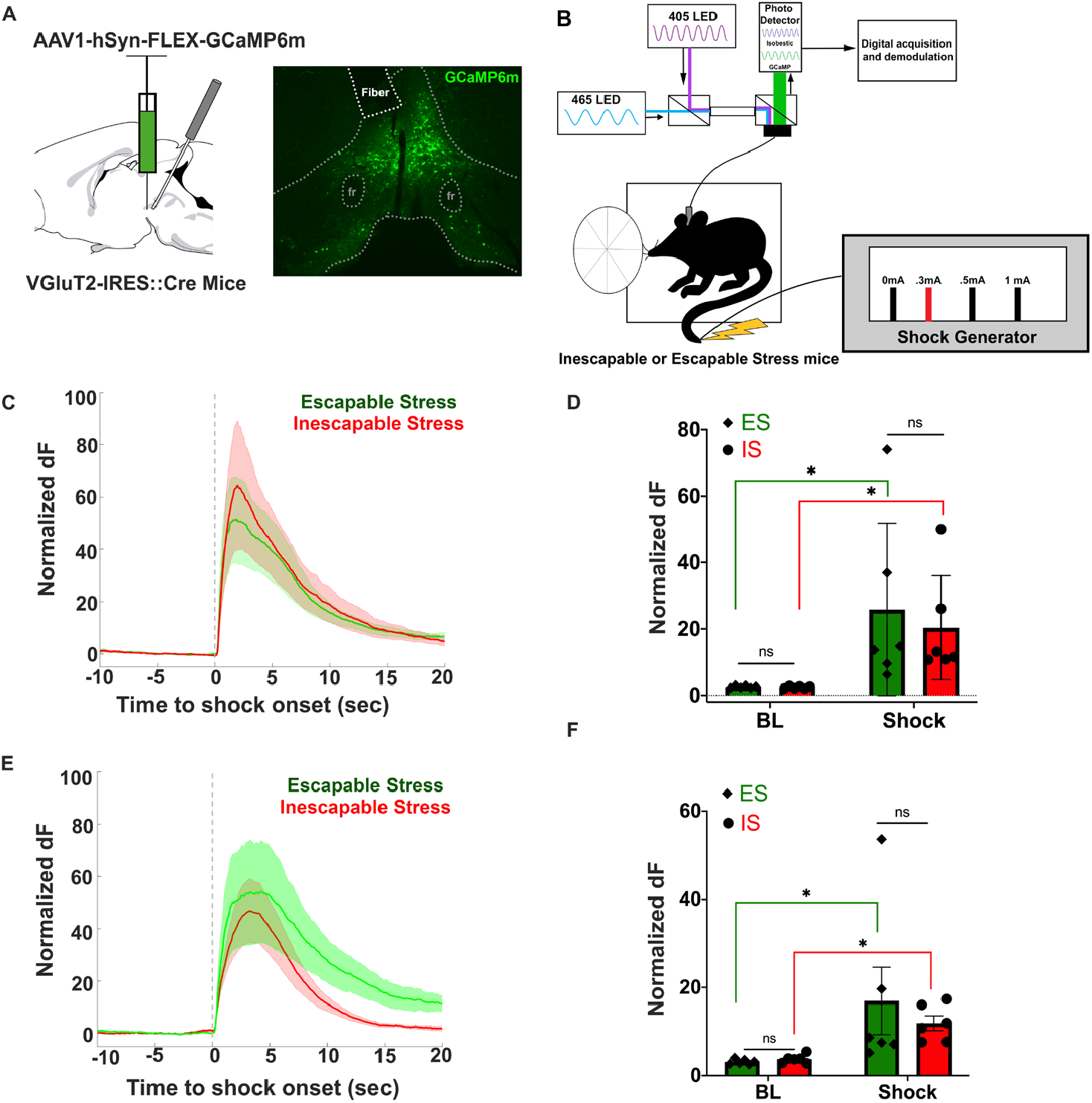
VTA glutamate neuron signaling is enhanced by both inescapable and escapable stress. (A) Schematic of viral delivery of Cre-inducible GCaMP6m and optical fiber implantation in VTA of VGluT2-IRES::Cre mice (left). Representative image of fiber placement and GCaMP6m expression within the medial VTA (right). fr = fasciculus retroflexus. (B) Integration of fiber photometry with the stressor controllability paradigm. Light path for fluorescence excitation and emission is through a single 400-µm optical fiber patch cord connected at the skull surface to the VTA fiber implant. (C) Mean trial-by-trial time course of average GCaMP6m z-scores event-locked to tailshock delivery and average peak fluorescence (D) over the first twenty trials. Dark red/green lines show mean fluorescence, while lighter shaded lines indicate standard error. (D) Individual normalized z scored data during the baseline (BL) period prior to shock and following tailshock (shock). Average fluorescence significantly increased compared to baseline for both escapable (ES) and inescapable (IS) mice, * p < 0.05. ES and IS did not differ between BL or Shock. (E) Mean trial-by-trial time course of average GCaMP6m z-scores event-locked to tailshock delivery and average peak fluorescence (F) over the final twenty trials. Dark red/green lines show mean fluorescence, while lighter shaded lines indicate standard error. (F) Average fluorescence significantly increased during tailshock compared to BL for both ES and IS, * p < 0.05, but they did not differ between groups. Data are mean ± SEM.

### Chemogenetic Inactivation of VTA Glutamate Neurons Prevents the Transsituational Outcomes of Uncontrollable Stress

Given that VTA glutamate neurons are sensitive to repeated trials of tailshock, we hypothesized that reducing the activity of this VTA cell population would mitigate the behavioral consequences of uncontrollable stress. In addition to sociability, we examined another transsituational outcome that follows uncontrollable stress in rats – enhanced shock-elicited freezing. We first validated a chemogenetic approach for reducing VTA glutamate activity using the inhibitory Gi-coupled designer receptor, hM4Di, and its actuator compound, low-dose clozapine (Gomez et al., 2017). VGluT2::Cre mice were injected with a Cre-dependent adeno-associated viral vector encoding hM4Di fused to a mCherry reporter and whole-cell recordings of VTA mCherry-expressing neurons were performed in the presence or absence of clozapine. Clozapine (10 µM) activation of the hM4Di receptor on VTA glutamate neurons significantly reduced the percent of current-evoked action potentials compared to baseline current-evoked action potentials, t(6) = 2.553, p = 0.043 (Figure 4A-C), demonstrating hM4Di-induced reduction of VTA VGluT2 firing.

**Figure 4.**
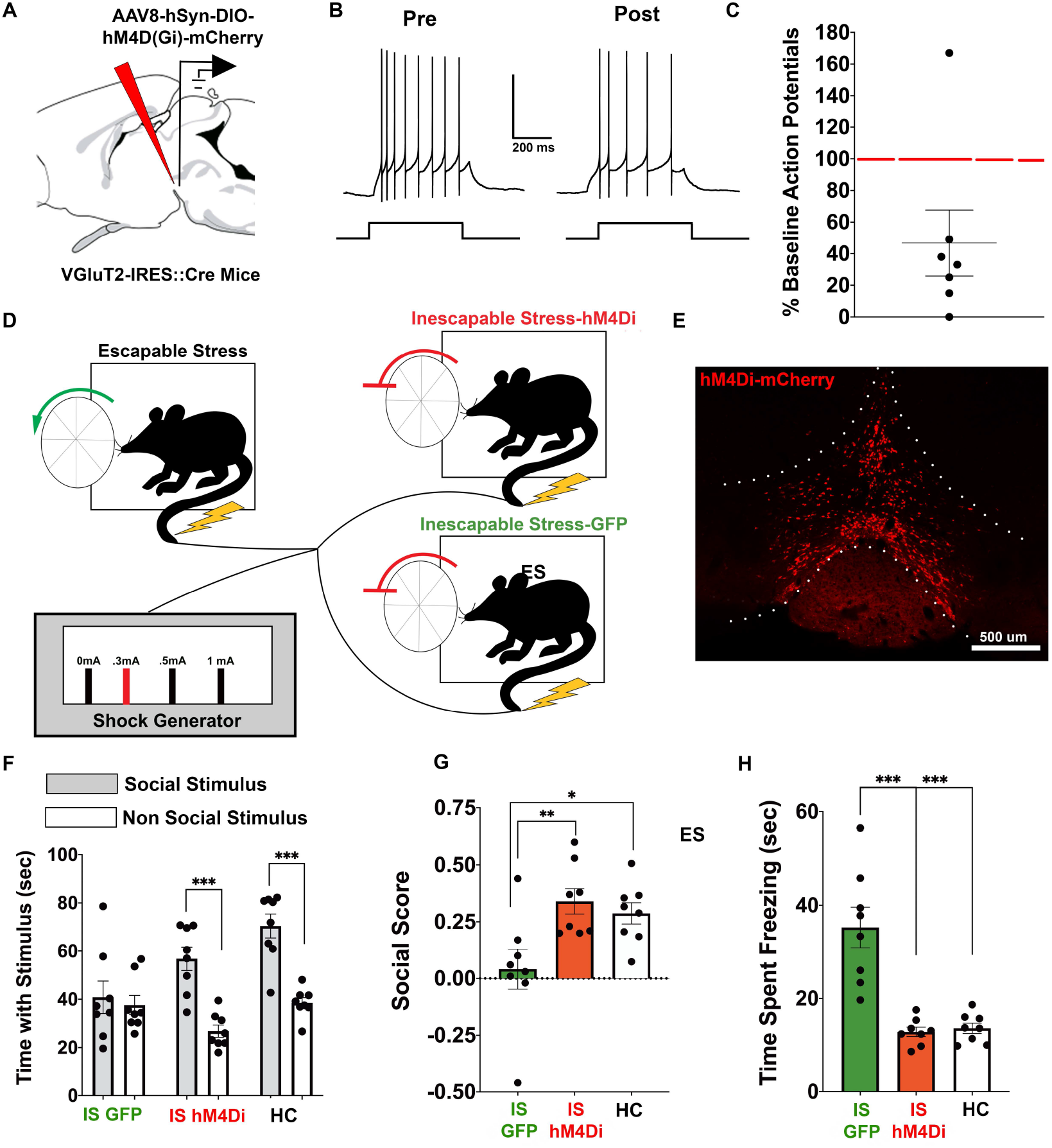
Chemogenetic suppression of VTA glutamate activity blocks the nonassociative behavioral consequences of inescapable stress. (A) Schematic of targeting Cre-inducible viral vector encoding the inhibitory designer receptor hM4D(Gi) to VTA in VGluT2-IRES::Cre mice. (B, C) Representative traces (B) and summarized data (C) from whole-cell recordings showing that the designer receptor actuator clozapine (10 µM) decreased the number of evoked action potentials in hM-4D(Gi)-mCherry VTA glutamate neurons. Red line indicates baseline firing. (D) Illustration of the experimental setup in which VGluT2::Cre male mice expressing Cre-dependent hM4Di-mCherry or GFP in VTA received yoked-inescapable shock (IS; 100 tail shocks, average intertrial interval 60 sec). Thirty minutes prior to IS or home cage treatment, IS hM4Di, IS GFP, and no stress home cage controls (HC) received clozapine (0.1 mg/kg; i.p.). (E) Representative image of hM-4Di-mCherry bilateral expression in the VTA of a VGluT2::Cre mouse. (F) Quantification of interaction behavior with a social (same-sex conspecific) or a non social (empty chamber) stimulus 24 hr following stress treatment. Home cage and IS hM4Di mice spent significantly more time with the social stimulus compared to the non social stimulus. IS GFP mice spent equal time with both social an non social stimulus types. (G) Preference for the social stimulus compared to the non social stimulus (social score) significantly differed between IS hM4Di and IS GFP groups. IS hM4Di and HC groups did not differ from one another. (H) Mean time spent freezing in response to the delivery of two footshocks in a novel context. IS GFP mice showed enhanced freezing which was absent in IS hM4Di and unstressed HC mice. Data are mean ± SEM. * p < 0.05, ** p < 0.01, *** p < 0.001; ANOVA with Sidak’s multiple comparison post hoc test.

To test the role of VTA glutamate neurons in mediating the behavioral outcomes of IS, VGluT2::Cre mice were injected in VTA with a Cre-dependent adeno-associated viral vector encoding hM4Di-mCherry (IS hM4Di mice) or a Cre-dependent control vector encoding GFP (IS-GFP mice or HC no stress mice). Uninjected VGluT2::Cre mice were assigned to ES in order to provide a pattern and duration of tailshock that was comparable to the fiber photometry experiment (Figure 4D, E). The ES mouse simultaneously controlled the duration, intensity, and temporal pattern of tailshocks for both yoked inescapable stress hM4Di mice and yoked inescapable stress GFP mice. Ten minutes prior to stress treatment, IS hM4Di and IS GFP mice were injected with a behaviorally subthreshold dose of clozapine (0.1 mg/ kg) (Gomez et al., 2017). A separate group of GFP-expressing mice were designated as no stress HC and were injected with the same dose of clozapine before being returned to the colony. ES mice acquired the instrumental controlling response, as demonstrated by a significant increase in the number of criterion quarter wheel-turns to terminate tailshocks over the shock session, Friedman’s test Χ2(9) = 17.03, p < 0.05. Sidak-adjusted pairwise comparisons showed that the criterion quarter wheel-turns completed during the initial block of ten tailshock trials was significantly less than the last block of ten tailshock trials, p < 0.01.

Twenty-four hours later, mice were assessed for social exploration followed by shock-elicited freezing. For the three-chamber sociability test, a mixed ANOVA yielded a Group x Stimulus interaction, F(2, 21) = 5.604, p < 0.05. Sidak-adjusted pairwise comparisons showed that IS hM4Di (p < 0.001) and HC (p < 0.001) groups spent significantly more time with the social stimulus than the non-social stimulus, but this social preference was absent in IS GFP mice (p > 0.05) **(Figure 4F)**. In addition, no stress HC mice spent significantly more time with the social stimulus than IS GFP mice (p < 0.01), but were not statistically different from IS hM4Di mice (p > 0.05). Further, IS hM4Di mice spent significantly less time with the non-social stimulus than HC mice (p < 0.05), but IS hM4Di mice did not differ from IS GFP mice in time with the non-social stimulus (p = 0.055). Thus, chemogenetic inhibition of VTA glutamate neurons during uncontrollable stress exposure blocks the social deficits that follow IS. In support, the social discrimination score was significantly lower in IS GFP mice compared to both IS hM4Di and HC groups (ANOVA: F(2, 21) = 6.291, p < 0.01; Sidak-adjusted pairwise comparisons, p < 0.05) **(Figure 4G)**.

Next, mice were assessed for shock-elicited freezing in a novel context and procedure room. During the two minute baseline exploration period, subjects showed minimal to no freezing in the novel chamber, suggesting that subjects did not generalize fear across the tailshock and footshock contexts, F(2, 21) = 2.229, p > 0.05. Total freezing was calculated over the two minutes following the first footshock. The univariate ANOVA yielded a significant main effect of Group, F(2, 21) = 26.153, p < 0.001. IS hM4Di and HC mice spent significantly less time freezing than IS GFP mice, all p < 0.001, indicating that silencing VTA glutamate neuronal activity during inescapable stress prevents subsequent IS-induced enhanced fear **(Figure 4H)**. Importantly, IS hM4Di mice and HC groups did not differ from one another in total time spent freezing, p > 0.05.

Together, these data indicate that VTA glutamate neuron activity during IS exposure is required for the social and fear-related transsituational outcomes of IS.

## DISCUSSION

Traumatic experiences can lead to a broad range of negative consequences that are supported by both associative (avoidance of or distress due to stimuli related to the trauma) and nonassociative (exaggerated transsituational responding to circumstances unrelated to the trauma) processes (Garcia-Keller et al., 2016; Lissek and Grillon, 2012; Lissek and van Meurs, 2015; Markowitz and Fanselow, 2020). Here, we adapted a stressor controllability paradigm from rats to mice in order to test the hypothesis that VTA glutamate neurons play a role in the nonassociative consequences of inescapable stress.

We first tested whether uncontrollable tailshock results in social deficits, a common feature of anxiety disorders (Kessler et al., 1999; Shin and Liberzon, 2010). Tailshock was explicitly chosen as an aversive stimulus over footshock because tailshock ensures that all subjects receive equivalent shock (duration, temporal pattern, intensity) that cannot be mitigated by postural adjustments or other behaviors such as jumping or locomotion (Maier, 2015). We found that inescapable stress results in reduced social exploration compared with unstressed mice. This social exploration deficit was observed in male but not female mice. While previous work has demonstrated differences in stress responses between sexes (Baratta et al., 2018), we believe that our protocol adapting stressor controllability from rats to mice is not yet optimized in female mice, rather than demonstrating an overt sex difference in stress responses. Just as recent research has optimized social defeat in female mice (Harris et al., 2018; Takahashi et al., 2017), future research will be needed to optomize stressor controllability in female mice.

Having identified that repeated tailshock stress results in social deficits in male mice, we next tested whether these social deficits were dependent on its uncontrollability. Mice that were allowed to escape tailshock stress showed no social deficits, similar to unstressed control mice, while mice that received identical tailshock stress again showed a social deficit involving a lack of discrimination between social and nonsocial stimuli. We interpret these results as demonstrating behavioral control over stress confers transsituational resilience against an inescapable stress-induced reduction in social exploration. Further, these results establish the stressor controllability paradigm in mice that have largely been performed in rats (Amat et al., 2010; Bai et al., 2017).

Following the identification that social deficits result from stressor uncontrollability, we examined the role of VTA glutamate neurons in stressor uncontrollability. Glutamate neurons within the VTA, defined by the expression of the vesicular glutamate transporter 2, contribute to reward, aversion, and threat (Barbano et al., 2020; Root et al., 2018). Optogenetic stimulation of VTA VGluT2+ neurons pathways to lateral habenula or nucleus accumbens shell results in place aversion (Qi et al., 2016; Root et al., 2014). Therefore, we performed two examinations: one investigating the extent to which VTA glutamate neuron activity is sensitive to stressor controllability and whether VTA glutamate neuron activity is required for the consequences of uncontrollable stress. Using GCaMP6m targeted to VGluT2+ cells within the VTA, we recorded population level changes in intracellular calcium during either escapable or inescapable stress. We hypothesized that when ES mice learned to terminate tailshocks by wheel turning, that this element of behavioral controllability would reduce VTA glutamate neuronal activity. Instead, we found that both ES and IS significantly increased intracellular calcium at tailshock onset and that there was no difference in calcium signaling when comparing their activity between ES and IS animals either during the first 20 trials or last 20 trials, where it might be expected that wheel turn escape responses would be fully associated with shock termination. Although there was no population level difference in VTA glutamate calcium signaling between the two stress groups, it is still possible that distinct VTA glutamate neurons differentially contribute to stressor control or loss of control. VTA glutamate neurons show heterogeneous electrophysiological dynamics for rewarding and aversive stimuli (Root et al., 2018) and receive distinct reward or aversion-related glutamatergic inputs (McGovern et al., 2020). Further, subtypes of VTA glutamate neurons defined by multiple genetic characteristics show different signaling patterns related to reward, aversion, and learned cues that predict these events (Root et al., 2020). Together, it is possible that distinct VTA glutamatergic cells or related circuits signal behavioral control or a lack of behavioral control. Single-cell approaches may better identify such task-related neurons. Nevertheless, these results provide further support that VTA glutamate neurons are highly sensitive to aversive events.

To determine whether VTA glutamate activity is necessary for the social deficits that result from inescapable stress we implemented a DREADD approach (Mahler et al., 2019). We expressed the inhibitory hM4Di DREADD to VGluT2+ cells within the VTA and administered a behaviorally-subthreshold dose of clozapine (Gomez et al., 2017) to selectively silence neuronal activity within these cells during inescapable stress. Whereas GFP-expressing mice showed social deficits following inescapable stress, chemogenetic silencing of VTA glutamate neurons prevented social exploration deficits following inescapable stress. hM4Di mice had comparable social scores as unstressed mice, both of which did not show the social deficits observed in GFP-expressing inescapable stress mice.

In addition to measuring social exploration following inescapable stress, we measured whether VTA glutamate neuron inactivation protects against exaggerated fear that can follow trauma (Asok et al., 2018; Kimble et al., 2014). Subsequent to social testing, we measured footshock-elicited freezing in a novel context and procedure room. Both unstressed and hM4Di mice showed low levels of freezing compared with the elevated levels of freezing observed in GFP mice. Thus, chemogenetic silencing of VTA glutamate neurons during inescapable stress blocked the development of exaggerated fear responses. There was no difference in freezing behavior prior to the first footshock between groups, highlighting the nonassociative aspects of this task by which associative fear from the wheel-turn box did not transfer to the footshock apparatus. Taken together with the results of the social test, the development of both social deficits and exaggerated fear responses by inescapable stress are blocked by chemogenetic inhibition of VTA glutamate neurons during inescapable stress.

Understanding which neural circuits and cell-types contribute to nonassociative deficits following stress is the first step in developing targeted therapeutics to ameliorate trauma-linked disorders. These experiments establish a mouse model of inescapable versus escapable stress, which can be leveraged with viral neuronal strategies such as optogenetics and neuronal recordings to tease apart stress neural circuitries. Additionally, this model provides a means to interrogate nonassociative deficits following stress, such as social exploration and enhanced fear. Using this model, we established that VTA glutamate neurons signal both escapable and inescapable stress and that silencing these neurons during inescapable stress prevents trans-situational stress-induced social exploration deficits and elevated fear. Additional work will be needed to determine which circuits and cellular subtypes VTA glutamate neurons are integrated in inescapable or escapable stress. VTA glutamate neurons are embedded within stress-related circuits, including the nucleus accumbens shell, lateral habenula, dorsal raphe, and lateral hypothalamus (Barbano et al., 2020; Dolzani et al., 2016; Faget et al., 2016; Lazaridis et al., 2019; Li et al., 2011; Qi et al., 2016; Root et al., 2014; Shumake and Gonzalez-Lima, 2003). This work sets a foundation for mechanistically exploring how VTA glutamate neurons and related neural circuitry confers stress susceptibility or resilience.

## ACKNOWLEDGEMENTS AND DISCLOSUREES

This research was supported by the Webb-Waring Biomedical Research Award from the Boettcher Foundation (DHR), R01 DA047443 (DHR), a 2020 NARSAD Young Investigator grant from the Brain and Behavior Research Foundation (DHR), R01 MH050479 (MVB), R21 MH116353 (MVB), American Australian Association Fellowship (MVB), and The University of Colorado. The funders had no role in study design, data collection and analysis, decision to publish, or preparation of the manuscript.

The authors report no financial interests or conflicts of interest.

## METHODS AND MATERIALS

### Animals

Male and female C57BL/6J and male Vglut2-ires-Cre knock-in mice (Slc17a6tm2(cre)Lowl/J; Stock #016963) were purchased from The Jackson Laboratory (Bar Harbor, ME) and bred at the University of Colorado. Mice were maintained in a colony room with a 12-hr light/dark cycle (lights on at 7:00 h), and with access to food and water ad libitum. All animal procedures were performed in accordance with the National Institutes of Health Guide for the Care and Use of Laboratory Animals and approved by the University of Colorado Boulder Institutional Animal Care and Use Committee.

### Inescapable stress

Male and female C57BL/6J mice (10-16 weeks) were randomly assigned to a single session of inescapable stress (IS) or no stress (home cage control; HC). IS mice were placed in a Plexiglas box (7 x 6 x 9 cm; Med Associates, St. Albans, VT) with a Plexiglas rod protruding from the rear of the box. The tail was secured to the rod with tape and affixed with copper electrodes and electrode paste. The session consisted of 100 tailshock trials (0.3 mA, 5 s duration each) on a variable interval 60-s schedule. Subjects were returned to the colony immediately following the tail shock procedure.

### Wheel-turn escapable (ES)/yoked IS procedure

For the manipulation of controllability, male C57BL/6J mice (10-16 weeks) were run in a triad design. Each subject of the triad was randomly assigned to a single session of escapable stress (ES), yoked-inescapable stress (Y-IS), or no tailshock stress (homecage, HC). ES and Y-IS mice were placed in separate Plexiglas boxes with a wheel mounted in the front (Med Associates). The tail was secured and affixed with two copper electrodes and electrode paste (as described above in Inescapable Shock). Each tailshock session consisted of 100 trials of tailshock (0.3 mA) on a variable interval 60-s schedule. Initially, the shock was terminated by a quarter turn of the wheel by the ES subject. When trials were completed in less than 7.5 s, the response requirement was in-creased by one step, to a maximum of three full turns of the wheel (i.e. twelve quarter-turns). Steps were 1, 2, 4, 8, and 12 quarter-wheel turns. The response requirement was reduced a step if the trial was completed between 7.5 and 25 s. If the requirement was not reached in less than 25 s, the shock was terminated at 30 s and the requirement was reduced to the initial one-quarter wheel-turn. For Y-IS subjects, the onset and offset of each tailshock was identical to that of ES. Thus, Y-IS subjects received identical shock intensity, duration, pattern, and number of shocks as the ES subject. During the designer receptor (DREADD) experiment, one ES mouse controlled tailshock for two Y-IS mice. The first Y-IS mouse was previously injected in VTA with a Cre-dependent GFP vector (IS GFP mice). These mice were controls to establish that neither the surgery nor the cellular production of a non-endogenous protein in these neurons was the driving force behind the behavioral effect. The second Y-IS mouse was previously injected in VTA with a Cre-dependent inhibitory designer receptor (hM4Di) tethered to mCherry (IS hM4Di mice). Both IS GFP and IS hM4Di mice were injected with a behaviorally subthreshold dose of clozapine (0.1 mg/kg) ten minutes prior to stress (Gomez et al., 2017). A separate group of mice previously injected in VTA with a Cre-dependent GFP vector were designated as no stress HC mice and were injected with the same dose of clozapine before being returned to the colony.

### Three-chamber sociability paradigm

Twenty-four hours after tailshock, social interaction levels were assessed using the three-chamber sociability assay. The test consisted of placing ES, IS, or HC subject in the center connecting compartment of an arena with three interconnected compartments (ANY-box, Stoelting, Wood Dale, IL). In one side, an empty sociability cage (Stoelting) was placed in the center and the other side contained an identical sociability chamber with a novel same-sex conspecific. The caged social mouse was approximately 2 weeks younger than the test mouse. Video recordings were made of mouse exploration of the entire arena for 10 min (30 Hz, ANY-maze, Stoelting). The social exploration test was quantified by counting the time the mouse spent exploring the cages containing the social and non-social stimuli. Exploration was defined as direct contact with the cage, including sniffing and rearing onto the apparatus. Social scores were quantified by the following formula: (social interaction time – nonsocial interaction time) / (social interaction time + nonsocial interaction time). Scores closer to 1 indicate a more social behavioral phenotype and scores closer to 0 reflect ambivalence.

### Shock-elicited freezing

Thirty minutes after the sociability paradigm, mice were brought to a separate novel room and placed inside a novel chamber with stainless-steel rod flooring (Med Associates). The shape, size, flooring, and illumination of the chamber differed from the context in which prior tailshock occurred. Subjects were allowed to explore the chamber for 2 min before receiving two scrambled footshocks (0.5 s, 0.5 mA, 1-min interstimulus interval) delivered through the grid floor. Within each chamber a video camera (10 Hz, Tucker-Davis Technologies, Alachua, FL) was connected to a computer for offline analysis of freezing behavior time-locked to shock onset (Synapse software, Tucker-Davis Technologies). Freezing was defined as the absence of movement except that required for respiration (Fanselow, 1980).

### Stereotactic surgery

Male VGluT2::Cre mice were anesthetized with 1-3% Isoflurane and secured in a stereotactic frame (Kopf). AAV8-hSyn-DIO-hM4Di-mCherry (Addgene, Cambridge, MA), AAV5-hSyn-DIO-GFP (Addgene), or AAV1-Syn-FLEX-GCaMP6m (Addgene) was injected into VTA (AP: -3.2 mm relative to bregma; ML: 0.0 mm relative to midline; DV: -4.3 mm from skull surface). All AAVs were injected at 2-5 x 10^12^ titer The total injection volume (500 nL) and flow rate (100 nL/min) were controlled with a microinjection pump (Micro4; World Precision Instruments, Sarasota, FL). Following injection, the needle was left in place for an additional 10 min to allow for virus diffusion, after which the needle was slowly withdrawn. For fiber photometry experiments, mice were additionally implanted with an optic fiber cannula (400 µm core diameter, 0.66 NA; Doric Lenses, Quebec, Canada) dorsal to VTA (AP: -3.2 mm relative to bregma; ML: -1.0 mm at 9.5° relative to midline; DV: -4.2 mm from skull surface) that was secured with screws and dental cement to the skull. All mice were allowed to recover for 3-4 weeks before experimentation.

### Whole-cell recordings

In order to confirm functionality of the inhibitory DREADD hM4Di in VTA glutamate neurons, whole-cell current clamp recordings were performed from VTA cells expressing the mCherry fluorophore fused to the hM4Di receptor in male VGluT2::Cre mice. Acute VTA slices were prepared from mice induced into a deep anesthetic plane with isoflurane and then rapidly decapitated. Brains were then removed and placed into ice-cold artificial cerebrospinal fluid (aCSF) containing the following (in mM): 126 NaCl, 2.5 KCL, 1.2 MgCl2^6H20, 2.4 CaCl2^H2O, 1.2 NaH2PO4, 11.1 glucose, and 21.4 NaHCO3, and bubbled with 95% O2 and 5% CO2. Coronal slices containing the VTA were cut at a thickness of 240 μm using a Leica VT1200S vibratome. Brain slices re-covered for at least 1 h at 37°C in aCSF containing the NMDA receptor antagonist MK-801 (15 μM). Slices were then transferred to a recording chamber and perfused with 37°C oxygenated aCSF at a flow rate of ∼2ml/min. Recording electrodes were made using a Narishige PC-10 vertical pipette puller. An internal recording solution containing (in mM): 135 potassium gluconate, 5 NaCl, 2MgCl2, 10 HEPES, 0.6 EGTA, 4 ATP, 0.4 GTP (pH 7.3) was used, and electrodes had a resistance of 1.6-2.2 MΩ when filled with this solution. Whole cell access was gained in voltage clamp (-60mV) and then switched to current clamp. Current clamp recordings were made at the cell’s resting membrane potential using an AxoPatch 200B amplifier (Molecular Devices, San Jose, CA). Electrophysiological data were collected at 10 kHz and filtered at 5 kHz using AxoGraph X software running on a MAC OS X operating system. Cells were periodically switched to voltage clamp to ensure that access and series resistance were stable throughout the recordings. Recordings were excluded from analysis if series resistance exceeded 18 MΩ or changed significantly over the course of the experiment. VTA glutamate cells did not fire spontaneous action potentials under these conditions. Thus, 600 ms, 30 pA current injections were applied at an interstimulus interval of 30 sec. Baseline firing in response to current injections was collected for 5 min followed by a 10 min bath application of 10 μM clozapine (Cayman Chemical, Ann Arbor, MI). The average number of action potentials generated during five consecutive sweeps was used to quantify baseline activity and the aver-age of five consecutive sweeps during clozapine application was used to quantify firing during hM4Di receptor activation. Due to the variance in action potential firing at baseline, the effect of clozapine was determined by normalizing to baseline.

### Calcium imaging

At least four weeks following intra-VTA viral injection, GCaMP6m recordings were performed in VGluT2::Cre mice during ES or Y-IS. GCaMP6m was excited at two wavelengths (465 nm and 405 nm isosbestic control) with amplitude-modulated signals from two light-emitting diodes reflected off dichroic mirrors and then coupled into an optic fiber (Barker et al., 2017; McGovern et al., 2020). Signals from GCaMP and the isosbestic control channel were returned through the same optic fiber and acquired with a femtowatt photo-receiver (Newport, Irvine, CA), digitized at 1kHz, and then recorded by a real-time signal processor (Tucker Davis Technologies). Behavioral timestamps of quarter-wheel turns and tailshock were digitized in Synapse by TTL input from a LabJack (Lakewood, CO). Analysis of the recorded calcium signal was performed using custom-written MATLAB scripts. Signals (465 nm and 405 nm) were downsampled (10X) and peri-event time histograms (PETHs) were created trial-by-trial between -20 sec and +30 sec surrounding each tailshock onset. For each trial, data was detrended by regressing the isosbestic control signal (405 nm) on the GCaMP signal (465 nm) and then generating a predicted 405 nm signal using the linear model generated during the regression. The predicted 405 nm channel was subtracted from the 465 nm signal to remove movement, photo-bleaching, and fiber bending artifacts (ΔF). Baseline normalized maximum z-scores were taken from -2 to 0 seconds prior to shock onset and maximum shock z-scores were taken from 0 to 2 seconds following tail shock.

### Histology

VGluT2::Cre mice were anesthetized with isoflurane and perfused transcardially with phosphate buffer followed by 4% (w/v) paraformaldehyde in 0.1 M phosphate buffer, pH 7.3. Brains were extracted, post-fixed overnight in the same fixative and cryoprotected in 18% sucrose in phosphate buffer at 4°C. Coronal sections containing the VTA (30 µm) were taken on a cryostat, mounted to gelatin-coated slides, and imaged for mCherry, GFP, or GCaMP6m expression and optical fiber cannula placement on a Zeiss Axioscope. Mice with optic fibers not localized to recording VTA GCaMP-expressing neurons were removed from the study.

### Statistics

Tests were conducted in SPSS (IBM) or Prism (GraphPad Software; San Diego, CA). For our initial examination of the effects of IS, a mixed 2 × 2 ANOVA tested the difference in time spent with a social and nonsocial stimulus between IS and HC groups. If the assumption of sphericity was not met (Mauchley’s test), the Greenhouse-Geisser correction was used. Sidak-adjusted pairwise comparisons followed up a significant interaction yielded from the ANOVA. A between-subjects t-test analyzed the difference in time spent freezing between stressed and control mice. Separate analyses were performed for each sex.

For the examination of stressor controllability, the measures for acquiring the wheel-turn controlling response were the mean criterion step achieved per ten trials as well as the mean latency to achieve criterion step per ten trials. Data were highly skewed and thus analyzed with Friedman’s nonparametric test. If the Friedman’s test yielded a significant effect, Sidak-adjusted simple pairwise comparisons tested the hypothesis that the first block of ten trials differed from other blocks of ten trials. A mixed 3 x 2 ANOVA tested the difference in time spent with a social and nonsocial stimulus between ES, Y-IS, and HC groups. A one-way ANOVAs tested the difference in social score between groups. A one-way ANOVA tested the difference in time spent freezing between groups. For all ANOVAs, if the assumption of sphericity was not met (Mauchley’s test), the Greenhouse-Geisser correction was used and Sidak-adjusted pairwise comparisons followed up a significant main effects or interactions. One outlier Y-IS mouse was excluded from the dataset in the controllability experiment. This mouse showed 5.94 standard deviations above the mean for time spent with the social stimulus.

In the DREADD experiment, time spent with the social stimulus and time spent with the non-social stimulus was compared between IS hM4Di, IS GFP, and no stress homecage groups using a mixed ANOVA. If the ANOVA yielded a significant group x stimulus interaction, Sidak-adjusted pairwise comparisons were used to compare between groups and stimuli. Time spent freezing was compared between the same groups using a univariate ANOVA. Pairwise comparisons followed up a significant main effect of the ANOVA.

Calcium imaging data was analyzed by comparing the average GCaMP z-score during baseline to the average z-score following shock using Wilcoxon sign ranked tests.

Electrophysiological data was analyzed by comparing the average number of action potentials generated in response to 5 consecutive 30 pA current injections during baseline and 5 consecutive 30 pA current injections during bath application of 10 uM clozapine. Data were normalized to baseline and a 1-sample t-test was used to determine if clozapine produced a significant change in the number of action potentials occurred in response to clozapine. 13

## Notes

### Competing Interest Statement

The authors have declared no competing interest.

